# Genome variants associated with RNA splicing variation in bovine are extensively shared between tissues

**DOI:** 10.1101/220251

**Authors:** Ruidong Xiang, Ben J. Hayes, Christy J. Vander Jagt, Iona M. MacLeod, Majid Khansefid, Phil J. Bowman, Zehu Yuan, Claire P. Prowse-Wilkins, Coralie M. Reich, Brett A. Mason, Josie B. Garner, Leah C. Marett, Yizhou Chen, Sunduimijid Bolormaa, Hans D. Daetwyler, Amanda J. Chamberlain, Michael E. Goddard

## Abstract

**Background:** Mammalian phenotypes are shaped by numerous genome variants, many of which may regulate gene transcription or RNA splicing. To identify variants with regulatory functions in cattle, an important economic and model species, we used sequence variants to map a type of expression quantitative trait loci (expression QTLs) that are associated with variations in the RNA splicing, i.e., sQTLs. To further the understanding of regulatory variants, sQTLs were compare with other two types of expression QTLs, 1) variants associated with variations in gene expression, i.e., geQTLs and 2) variants associated with variations in exon expression, i.e., eeQTLs, in different tissues.

**Results:** Using whole genome and RNA sequence data from four tissues of over 200 cattle, sQTLs identified using exon inclusion ratios were verified by matching their effects on adjacent intron excision ratios. sQTLs contained the highest percentage of variants that are within the intronic region of genes and contained the lowest percentage of variants that are within intergenic regions, compared to eeQTLs and geQTLs. Many geQTLs and sQTLs are also detected as eeQTLs. Many expression QTLs, including sQTLs, were significant in all four tissues and had a similar effect in each tissue. To verify such expression QTL sharing between tissues, variants surrounding (±1Mb) the exon or gene were used to build local genomic relationship matrices (LGRM) and estimated genetic correlations between tissues. For many exons, the splicing and expression level was determined by the same *cis* additive genetic variance in different tissues. Thus, an effective but simple-to-implement meta-analysis combining information from three tissues is introduced to increase power to detect and validate sQTLs. sQTLs and eeQTLs together were more enriched for variants associated with cattle complex traits, compared to geQTLs. Several putative causal mutations were identified, including an sQTL at Chr6:87392580 within the 5^th^ exon of kappa casein (*CSN3*) associated with milk production traits.

**Conclusions:** Using novel analytical approaches, we report the first identification of numerous bovine sQTLs which are extensively shared between multiple tissue types. The significant overlaps between bovine sQTLs and complex traits QTL highlight the contribution of regulatory mutations to phenotypic variations.

## Background

Cattle are an important source of meat and dairy products for humans worldwide. Also, cattle can be used as clinical models to study genetic causes of human diseases [1]. To improve productivity, health performance and efficiency of cattle, traditional selective breeding has been widely used. In the last decade, genomic selection, originally developed in cattle breeding, has further increased the rate of genetic improvement of complex traits in all livestock species [2, 3]. However, genomic selection commonly uses genotyping arrays that are based on single nucleotide polymorphisms (SNPs) of which very few have known biological functions or directly impact genetic variation in production traits. Knowledge of the genes involved and polymorphic sites would increase our understanding of the biology and may further increase the rate of genetic improvement [4].

Many of the sequence variants that are associated with complex traits (quantitative trait loci or QTL) are not coding variants and are presumed to influence the regulation of gene expression, that is to be expression QTLs [5]. An expression QTL might be associated with the variation in overall transcript abundance from the gene, which we will refer to as a gene expression QTL or geQTL. In cattle and humans, geQTLs show significant enrichments for mutations associated with diseases and complex traits [5–7].

After transcription, RNA is spliced by intron removal and exon ligation to create various mature transcripts. An expression QTL associated with the changes in the expression ratio of an exon to the gene implies that it alters RNA splicing. This type of expression QTL is then defined as a splicing QTL or sQTL, which have been studied in humans by inferring the individual splicing ratio from RNA sequence data [8]. More recently, sQTLs, identified using intron information extracted from RNA sequence data, were demonstrated to have fundamental links with human diseases [9, 10]. RNA splicing also results in different expression levels of exons within a gene. Thus, in theory, the type of expression QTL that change the level of expression of one or several exons, i.e., exon expression QTL or eeQTLs, may represent some sQTLs. However, the extent to which eeQTLs overlap with sQTLs and/or geQTLs remains unclear, at least in cattle.

Knowledge of large mammal regulatory mutations is limited mainly to humans, where there have been multiple studies reporting on expression QTLs [9, 11, 12]. In this study, we aim to identify bovine cis splicing QTLs using the abundances of genes, exons and introns estimated from RNA sequence data from hundreds of animals and multiple tissues along with imputed whole genome sequences. We examined the extent to which sQTLs can be detected in white blood cells, milk cells, liver and muscle transcriptomes and the extent to which sQTLs overlap with conventional QTL associated with complex traits overlapped. To further characterise the features of sQTLs, we used the counts of genes and exons to map another two types of cis expression QTLs: eeQTLs and geQTLs, and then analysed their relationships with sQTLs in different tissues.

## Results

Data quality

In total, we analysed 378 transcriptomes of 19 tissue types from 214 cattle generated from four experiments covering major dairy and beef cattle breeds (Table 1, Figure 1a and Additional file 1: Supplementary Methods). Following recommendations from Mazzoni and Kadarmideen 2016 [13] RNA sequence quality was assessed and was detailed in Additional file 1: Supplementary Methods. Based on the results produced by Qualimap 2 [14], no significant events of RNA degradation were observed in all studied tissues (Additional file 2: Supplementary Figure S1). Also, according to Qualimap 2 [14], on average, 60.3% of reads were mapped to exonic regions of the bovine reference genome (UMD3.1), 15.4% of reads were mapped to intronic regions and 24.4% were mapped to integenic regions (Additional file 3: Supplementary Table S1). Splicing junction annotation and saturation were estimated using RSeQC [15]. As a small demonstration, no significant difference was observed in the annotated splicing junction events in the bovine reference genome (UMD3.1), using different RNA-seq alignment software including HISAT2 [16], STAR [17] and TopHat2 [18] (Additional file 4: Supplementary Table S2). Although HISAT2 and STAR outperformed TopHat2 for novel splicing junction events (do not exist in the current bovine UMD3.1 genome). Also, using a splicing junction saturation analysis for all tissues we observed saturated coverage for the known splicing junctions (Additional file 2: Supplementary Figure S2), though there appeared to be more potential for splicing junction discovery for the novel category.

**Table 1.** Summary of experiments, data and analyses. Tissue splicing: variation in differential splicing associated with tissue types estimated using RNA sequence data from all experiments. Breed splicing: variation in differential splicing associated with Holstein and Jersey breeds estimated using RNA sequence data of milk cells from experiment III. sQTLs: cis splicing quantitative trait loci, sQTLs estimated using RNA sequence data and imputed whole genome and from experiment III and IV. Data from experiment III and IV were also used to estimate exon expression eeQTLs and gene expression geQTLs.

| Tissue splicing | Breed splicing | sQTLs | Experiment | Tissue type | Sample No. | Breed | Individual No. |
| --- | --- | --- | --- | --- | --- | --- | --- |
| ✓ |  |  | I | 18 various<sup>a</sup> | 54 | Holstein | 1 |
| ✓ |  |  | II | milk cells & mammary | 12 | Holstein | 6 |
| ✓ |  | ✓ | III | white blood cells | 105 | Holstein | 105<sup>b</sup> |
| ✓ | ✓ | ✓ | III | milk cells | 131 | Holstein & Jersey | 105<sup>b</sup> & 26 |
| ✓ |  | ✓ | IV | liver | 35 | Angus | 35 |
| ✓ |  | ✓ | IV | muscle | 41 | Angus | 41 |

<sup>a</sup>: 18 tissues from [24]. <sup>b</sup>: The same 105 Holstein cattle, each of which had both white blood and milk cell transcriptomic data.

**Figure 1.**
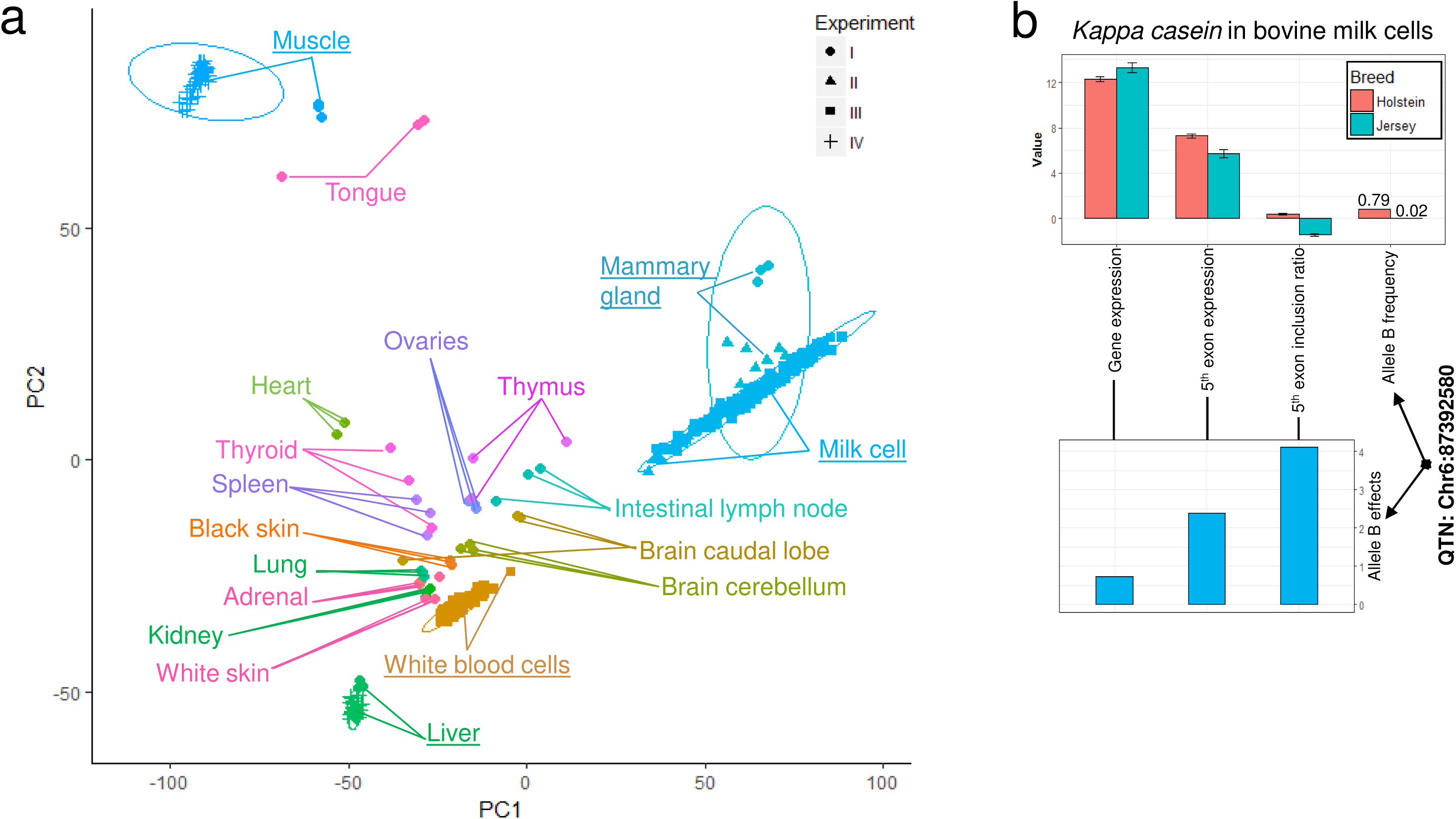
**a**: Sample principal components clustering based on exon expression. Circles on the plot were ellipses drawn based on tissue types at the confidence interval = 0.95. Tissue types with which the non-overlapping ellipses were drawn were emphasised with underscored text labelling. Ellipses that were drawn based on experiments can be found in Supplementary Figure S4. **b**: The significant splicing events between breeds and between genotypes (cis splicing quantitative trait loci, sQTLs) for *CSN3* in the milk cell transcriptome. In the upper panel, from left to right: the 1^st^ pair of bars are the least square means of normalised expression level of the gene (ENSBTAG00000039787) in Holstein and Jersey breeds; the 2^nd^ pair of bars are the normalised expression level of the 5^th^ exon (6:87392578-87392750) in Holstein and Jersey breeds; the 3^rd^ pair of bars are the normalised inclusion ratio of the 5^th^ exon in Holstein and Jersey breeds; and 4^th^ pair of bars is the frequency of the B allele of the sQTL (Chr6:87392580) for *CSN3* in Holstein and Jersey breeds. The standard errors bars are indicated. In the lower panel, from left to right: the 1^st^ bar is the effects (signed t values, b/se) of the sQTL (Chr6:87392580) B allele on the normalised expression of the gene; the 2^nd^ bar is the sQTL B allele effect on the normalised expression of the 5^th^ exon; and the 3^rd^ bar was the sQTL B allele effects on the normalised inclusion ratio of the 5^th^ exon.

Animals with white blood cell RNA-seq data were evaluated for the consistency between imputed genotypes from the 1000 bull genomes project [19] and RNA sequence genotypes as predicted from the RNA sequence data using samtools [20]. On average, the concordance between imputed sequence genotypes and RNA sequence genotypes was 0.943 (Additional file 2: Supplementary Figure S3), which was consistent with the average imputation accuracy (0.926) of the 1000 bull genomes project [21]. The comparison of the genotypes was detailed in Additional file 1: Supplementary Methods.

Overall, samples from the same or similar tissues clustered together rather than clustering by experiments, based on exon expression levels (Figure 1a). This was supported by further analyses of the clusters where ellipses, drawn based on tissue types, were clearly separated (Additional file 2: Supplementary Figure S4a,c,e), whereas ellipses drawn based on different experiments overlapped (Additional file 2: Supplementary Figure S4b,d,f), at the confidence interval = 0.95 [22]. Consistent with previous reports [23], milk cells and mammary gland transcriptomes were closely related.

### Differential splicing between tissues and breeds

We primarily defined a differentially spliced gene as a gene which contained exons whose inclusion ratios (exon expression divided by gene expression) were significantly associated with tissue or breed (FDR<0.1). To verify the significantly spliced exons, we imposed a requirement that at least one adjacent intron had an excision ratio [9, 25] that was also significantly (FDR<0.1) associated with tissue or breed (See methods). The FDR threshold of such exon splicing events was considered as approximately 0.01 by combining the FDR thresholds from exon and intron analyses (0.1 × 0.1). The overlaps of genes displaying differential splicing from exon and intron analyses were shown and examined in 2 × 2 tables by Chi-square tests (Additional file 5: Supplementary Table S3). Overall, the overlap of the results from exon and intron analyses was significantly more than expected by random chance.

Using data from all experiments (Table 1), there were 8,657 genes in which at least one exon had the variation in splicing associated with differences between tissue types. A list of these genes with the significances of differential splicing for the exons and introns was shown in Additional file 6: Supplementary Table S4. The top 10% of these significantly differentially spliced genes had a GO term enrichment (FDR<0.01) of ‘regulation of cellular process’, suggesting very general roles of these genes in cell function. There were148 genes with significant differential splicing events in the milk cell transcriptome between breeds (Table 1, Additional file 7: Supplementary Table S5). While these genes did not show any significant GO term enrichments, they included the milk protein gene *CSN3* [26, 27], where the 5^th^ exon (6:87,392,578-87,392,750) was more commonly included in the transcript in Holstein cattle than in Jersey cattle (Figure 1b).

### cis splicing quantitative trait loci (sQTLs)

The mapping of sQTLs was based on data from 312 transcriptomes generated from experiments III and IV, including white blood cells, milk cells, liver and muscle tissues (Table 1). In total 207 individuals had imputed whole genome sequence data and in experiment III, 105 genotyped cattle had both white blood and milk cell transcriptome data (Table 1). Similar to differential splicing analyses described above, a cis-acting sQTL was defined as a SNP significantly (FDR<0.1) associated with the variation in the inclusion ratio of the exon (up to 1Mb away) and significantly (FDR<0.1) associated with the variation in at least one adjacent introns’ excision ratio [9, 25]. When analysed separately, the overlap between sQTLs found by exon analyses and sQTLs found by intron analyses were significantly more than expected by random chance (Additional file 5: Supplementary Table S3). After requiring that the variation in the inclusion and excision ratios for adjacent exons and introns both be associated with the same SNP, 138,796 sQTLs were called in the white blood cells, 28,907 sQTLs were called in the milk cells, 11,544 sQTLs were called in the liver tissue and 5,783 sQTLs were called the muscle tissue (Figure 2, Additional file 5, 8: Supplementary Table S3 and S6).

**Figure 2.**
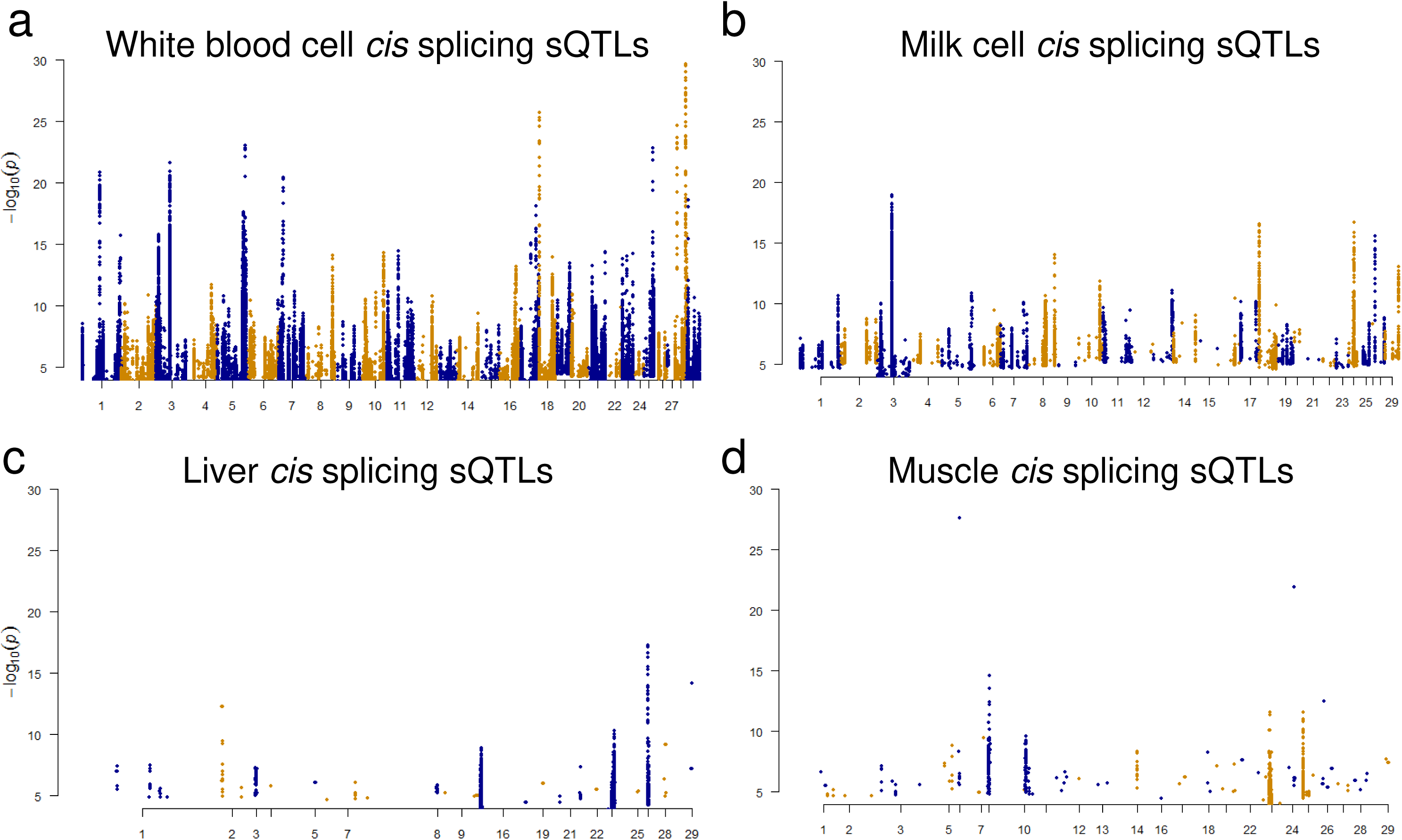
Manhattan plots of significant cis splicing quantitative trait loci (sQTLs, approximate FDR<0.01 and within 1Mb of the exon) in white blood cells (**a**), milk cells (**b**), liver tissue (**c**) and muscle tissue (**d**). A significant sQTLs was defined as a SNP associated with the variation in the exon inclusion ratio and also variation in at least one excision of an adjacent intron at the same significance level. The input SNPs had significance p<0.0001. sQTLs in all tissues with their associated genes and significance are given in Supplementary Table S6.

The significant sQTLs in white blood and milk cells were mapped to 929 and 283 genes, respectively (Table 2). Many SNPs were significant for sQTLs due to linkage disequilibrium between SNPs close to the same gene. The results do not imply many sQTLs per gene.

**Table 2.** Summary of expression QTLs detected. cis sQTLs: significant SNPs within ± 1Mb of the exon, associated with the variation in its inclusion ratio and also associated with the variations in the excision of an adjacent intron at the same significance level. Where the FDR threshold was approximated to 0.01 by combining FDR thresholds used in exon (FDR < 0.1) and intron (FDR < 0.1) analyses. cis eeQTLs: significant (FDR<0.01) SNPs within ± 1Mb of the exon, associated with the variation in its abundance. cis geQTLs: significant (FDR<0.01) SNPs within ± 1Mb of the gene associated with the variation in its abundance.

| cis QTL type | Tissue | Expression QTL number | Gene number |
| --- | --- | --- | --- |
| sQTLs | white blood cells | 138,796 | 929 |
|  | milk cells | 28,907 | 283 |
|  | liver | 11,544 | 49 |
|  | muscle | 5,783 | 76 |
| eeQTLs | white blood cells | 802,685 | 6,446 |
|  | milk cells | 100,844 | 2,102 |
|  | liver | 37,322 | 346 |
|  | muscle | 64,675 | 1,267 |
| geQTLs | white blood cells | 96,530 | 842 |
|  | milk cells | 4,099 | 99 |
|  | liver | 39,306 | 419 |
|  | muscle | 57,054 | 1,150 |

In the milk cell transcriptome, the fifth exon of *CSN3* (6:87392578-87392750), which as described above was differentially spliced in Holstein and Jersey cattle (Figure 1b), and was strongly associated with an sQTL (Chr6:87392580, p = 5.0e-07, Additional file 8: Supplementary Table S6). This sQTL is physically located within the 5^th^ exon of *CSN3*. Also, the B allele of this sQTL increased the expression and inclusion ratio of the 5^th^ exon and had a higher allele frequency among Holstein cattle than Jersey cattle (0.79 vs 0.02). This predicted that the expression and inclusion ratio of the 5^th^ exon would be significantly higher in the Holstein cattle than Jersey cattle, which was in line with the observations in Figure 1b. In addition, this sQTL was also predicted to be a splice site (‘splice_region_variant’) by Ensemble [28] and NGS-SNP software [29]. Much smaller numbers of significant sQTLs were detected in liver (11,544 SNPs) and muscle (5,783 SNPs) (Figure 2c,d). This was probably due to the smaller sample size (Table 1) and lower sequence depth of liver and muscle from experiment IV (Additional file 1: Supplementary Methods) than that of white blood and milk cells from the experiment III (Figure 2a,b, Table 2).

### Comparing sQTLs with exon expression eeQTLs and gene expression geQTLs

Many more significant eeQTLs than sQTLs were detected in all tissues studied (Table 2). In white blood and milk cells, the number of geQTL was smaller than the number of significant sQTLs in white blood and milk cells (Table 2).

Figure 3a showed that sQTLs were a median distance of about 200 kb from the transcription start site (TSS) and were slightly closer to the TSS than eeQTLs and geQTLs. All 3 classes of expression QTLs had a lower percentage of intergenic SNPs and a higher percentage of intronic and coding SNPs, including splice sites than the same categories across all SNPs analysed (Figure 3b). Specifically, sQTLs had the highest percentage of intronic SNPs, compared to eeQTLs and geQTLs. However, no consistent ranking of concentrations of ‘Splice’ SNP category for sQTLs, eeQTLs and geQTLs were observed in different tissues (Figure 3b).

**Figure 3.**
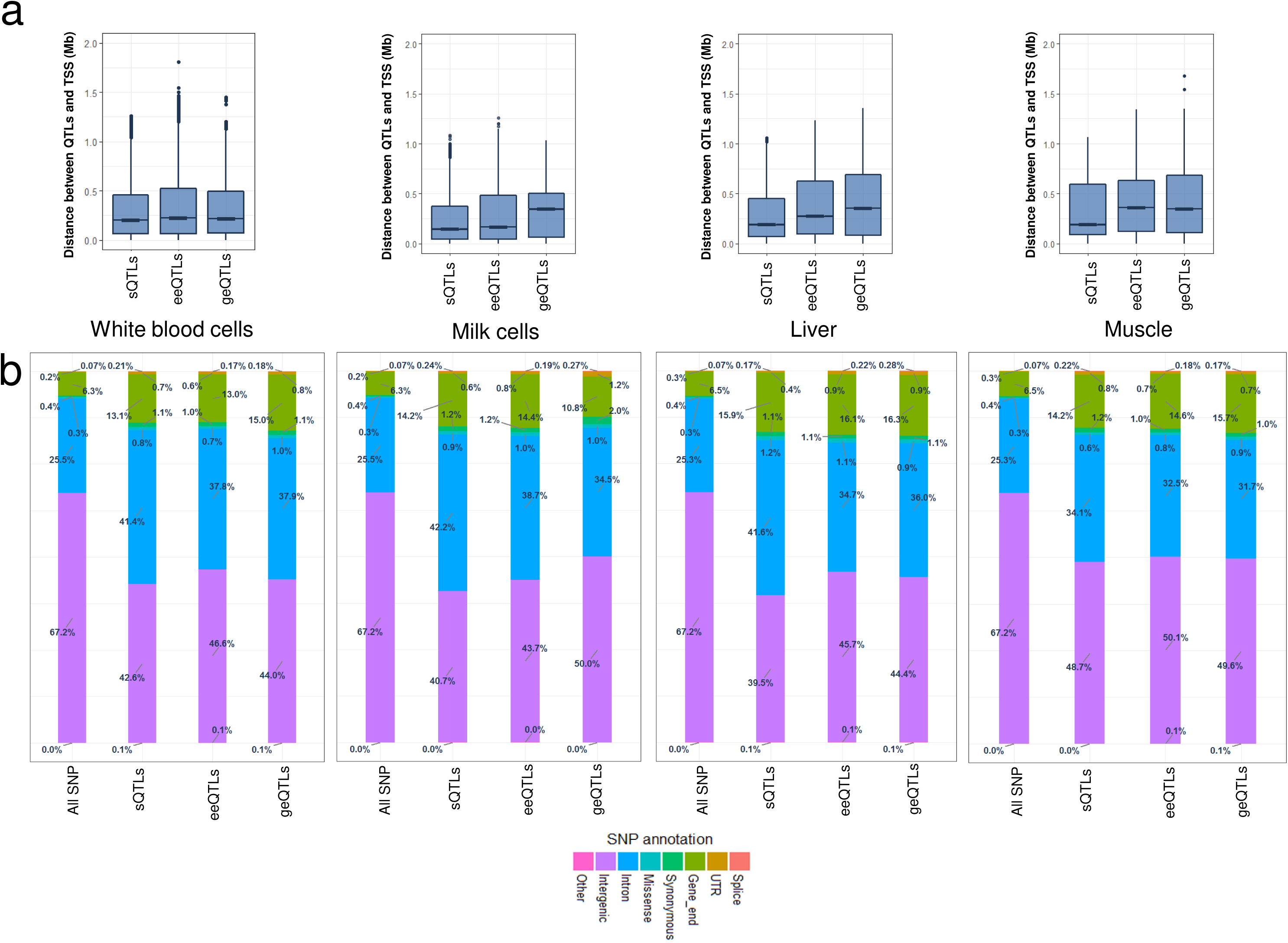
Features of cis splicing quantitative trait loci (sQTLs) compared to exon expression QTL (eeQTLs) and gene expression QTL (geQTLs). **a**: The distance between the transcription start site (TSS) and the expression QTLs. TSS information was downloaded from Ensembl (bovine reference UMD3.1). **b**: The proportion of expression QTLs annotated as splice, UTR, gene_end, synonymous, missense, intron, intergenic or other. SNP annotations were based on Variant Effect Predictor. ‘Splice’ included all SNP annotations containing the word ‘splice’. ‘UTR’ included 3’ and 5’ untranslated region. ‘Gene_end’ included upstream and downstream.

### Shared genetic influences between cis QTL types

Within each tissue, the sharing of SNPs between all three expression QTL types was significantly more than expected by chance (Figure 4). However, in the white blood and milk cells, which had relatively large sample size (n>=105, Table 1), the largest absolute amount of SNP sharing appeared to be between sQTLs and eeQTLs (Figure 4). This was followed by the amount of SNP sharing between eeQTLs and geQTLs (Figure 4a,b). In liver and muscle tissue which had relatively small sample size (n<=41) and low sequencing depth, the largest absolute amount of SNP sharing was between eeQTLs and geQTLs, followed by the amount of SNP sharing between sQTLs and eeQTLs (Figure 4c,d).

**Figure 4.**
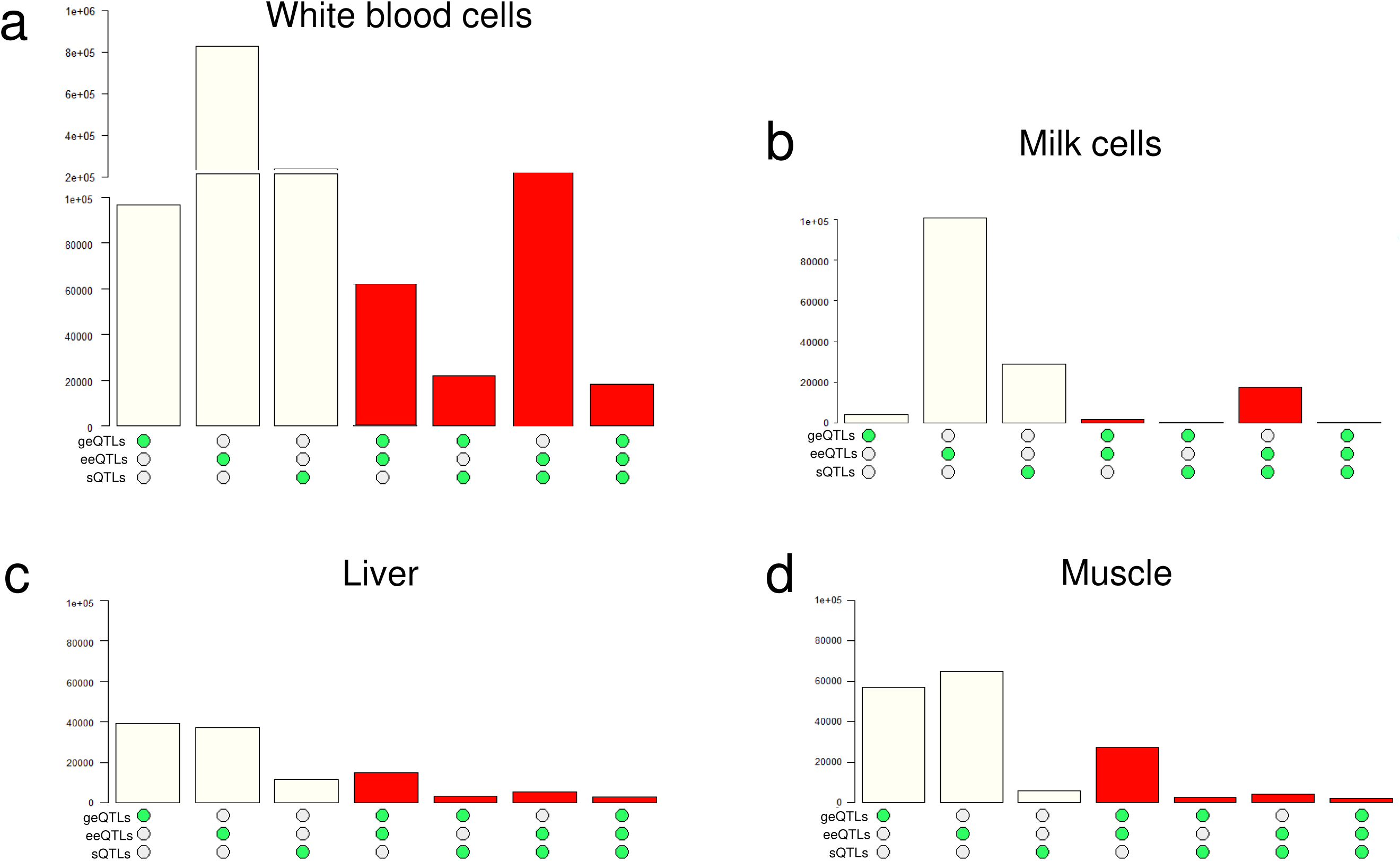
Overlaps of different expression QTL types for white blood cells (**a**), milk cells (**b**), liver(**c**) and muscle (**d**). Within each panel, y-axis was the number of significant expression QTLs; from left to right as guided by the green dots, the 1^st^ bar indicated the number of significant cis splicing QTL (sQTLs); the 2^nd^ bar indicated the number of significant exon expression QTL (eeQTLs); the 3^rd^ bar indicated the number of significant gene expression QTL (geQTLs); the 4^th^ bar indicated the number of SNPs identified as both geQTL and eeQTL; the 5^th^ bar indicated the number of SNPs identified as both geQTL and sQTL; the 6^th^ bar indicated the number of SNPs identified as both eeQTL and sQTL; and the 7^th^ bar indicated the number of SNPs identified as geQTL and eeQTL and sQTL. The red colour indicates that the overlap between categories of expression QTLs was significantly more than expected by random chance based on Fisher’s exact test.

To further examine the relationship between sQTLs and eeQTLs, a two by two table of sQTL and eeQTL counts in white blood and milk cells, which had comparable sample sizes, was created (Additional file 9: Supplementary Table S7). This suggested that when an sQTL was found, it was highly likely to be also identified as an eeQTL. For example, of 138,796 sQTLs found in the white blood cells, 109,155 of them were also blood eeQTLs, but only 21,766 of them were identified as blood geQTLs. Again, for these 138,796 blood sQTLs, although only 18,005 and 25,932 of them were milk sQTLs and eeQTLs, respectively, an even smaller number, 720, of them were identified as milk cell geQTLs.

### Shared genetic influences between tissues

Within each type of expression QTL between different tissues, the majority of the significant expression QTL sharing was observed between white blood and milk cells and between liver and muscle (Figure 5a). This is not unexpected since most of the white blood and milk cells came from the same lactating cows and the muscle and liver from different growing Angus bulls. Nevertheless, there was significant sharing of eeQTLs between milk cells and liver and between milk cells and muscle (Figure 5a). The largest amount of across-tissue expression QTL sharing was observed in eeQTLs, followed by sQTLs and geQTLs (Figure 5). Where a SNP was significantly associated with variation in expression in two tissues, the direction of effect was usually the same in both tissues (Additional file 2: Supplementary Figure S5). The correlation between effects of expression QTLs for white blood cells and milk cells (Additional file 2: Supplementary Figure S5a,c,e) was stronger than that between liver and muscle (Additional file 2: Supplementary Figure S5b,d,f). The sharing at the SNP level between white blood cells and milk cells and between liver and muscle were also evident at the exon and gene level (Figure 5b).

**Figure 5.**
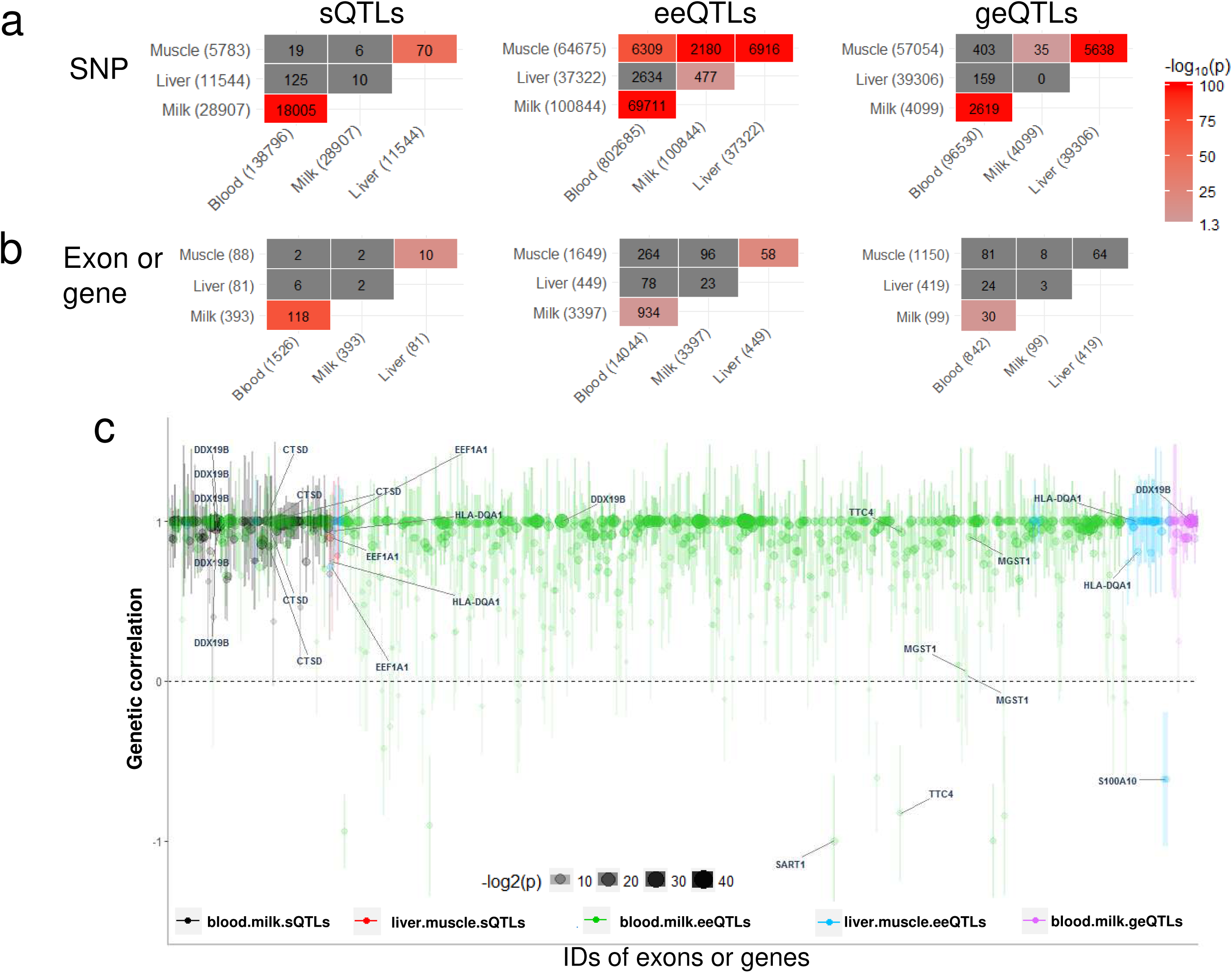
Shared genetic influence on the splicing, exon and gene expression between tissues. Blood refers to white blood cells and milk refers to milk cells. **a**: Each matrix shows the pair wise comparison of the numbers of significant SNP and the total number of significant SNPs detected for each analysis shown in parentheses. The significance of each overlap was tested by Fisher’ exact test, given the total number of SNP analysed and the total number of significant SNP, the result of which is represented by the colour of that position in the matrix. **b**: Each matrix shows the pair wise comparison of the numbers of exon/gene with significant associations and the total number of exon/gene detected with significant associations for each analysis shown in parentheses. In panel **b**, the numbers were either exon numbers for sQTLs (splicing quantitative trait loci) and eeQTLs (exon expression quantitative trait loci) or gene numbers for geQTLs (gene expression quantitative trait loci. **c**: Between tissue genetic correlations of either the inclusion ratio of the exons, the expression of the exons or the expression of the genes that had significant sharing of expression QTLs in panel **a.** Dot size and transparency were negatively correlated with p value of the significance of the genetic correlation being different from 0. The error bars of the genetic correlation were shown in vertical lines of each dot. Some genes of interests were highlighted.

The expression QTL sharing between tissues was further examined for all types of expression QTL by using a less stringent p-value (p<0.05) to test their effect (Additional file 2: Supplementary Figure S6). This showed that the expression QTL sharing between tissues was stronger for sQTLs and eeQTLs (Additional file 10: Supplementary Figure S6a-d), than the sharing for geQTLs (Additional file 10: Supplementary Figure S6e-f). Again, more expression QTL sharing was found between white blood cells and milk cells than between liver and muscle. For instance, 75% of eeQTLs significant in the white blood cells at p<0.002 were significant in milk cells at p<0.05 (Additional file 2: Supplementary Figure 6c).

The correlation between estimated SNP effects on gene splicing and expression in different tissues are lower in magnitude than the true correlation between SNP effects, because the effects are estimated with error and these errors are independent between tissues. To estimate the true correlation, we computed the genetic correlation between SNP effects in two different tissues by GREML^16^ using a local genomic relationship matrix or LGRM built from SNPs from 1 Mb surrounding the exon or gene (Figure 5c). LGRM differs from a conventional GRM by focusing on the local SNPs (in this case within 1 Mb distance) with potential *cis* genetic associations with the variation in the splicing or expression level of the exon or gene. This was in agreement with the definition of the *cis* expression QTLs which were also within 1Mb distance to the exon or gene in the current study. Out of 1,145 analysed sQTLs (inclusion ratio of the exon), eeQTLs (expression level of the exon) and geQTLs (expression level of the gene) between tissues, 598 had genetic correlations significantly (p<0.05) different from 0, out of which 561 had genetic correlations insignificantly (p >= 0.05) different from 1. That is, in many cases, the variation in exon expression in white blood cells and milk cells was associated with the same *cis* polymorphism (s).

Often both splicing events and exon expression within a gene were highly correlated between white blood and milk cells, for instance *DDX19B*, *CTSD* and *EFF1A1* (Figure 5c). In liver and muscle, exons from *HLA-DQA1* encoding major histocompatibility complexes [30] also showed significant genetic correlations between tissues based on both exon expression and splicing. There were more cases of eeQTLs than sQTLs and geQTLs and so there were more estimates of genetic correlations between white blood and milk cells in Figure 5c. The genetic correlations between eeQTLs in white blood and milk cells show a range from +1 to - 1 although most are close to +1. Exons with negative genetic correlations of expression between white blood cells and milk cells were mapped to *SART1* [30], a post-transcriptional regulator in epithelial tissues and *TTC4* with potential to mediate protein-protein interactions [30]. These negative genetic correlations imply that there are mutations that increase the expression of the exon in milk cells but decrease it in white blood cells. An exon within *S100A10*, a cell cycle progress regulator, showed negative genetic correlation of expression between liver and muscle.

Genetic correlations between exon expression levels in two tissues can be different between exons within the same gene. For example, only the 2^nd^ exon (5:93,942,055 −93,942,195) of *MGST1* (which is associated with the variation in dairy cattle milk fat yield [27, 31]) had a significant genetic correlation of expression between white blood cells and milk cells (Figure 5c). This was largely due to a few eeQTLs with relatively highly significant effects (p<1e-10) on the expression levels of the 2^nd^ exon in the milk cells and a similar but less significant effect (p<1e-4) on the 2^nd^ exon expression in white blood cells (Additional file 2: Supplementary Figure S7). For exons 1 and 3, the significances of the eeQTLs in both milk and white blood cells were > 1e-3. For exon 4, the significance of the majority of eeQTLs in both milk and white blood cells were > 1e-5 (Additional file 2: Supplementary Figure S7).

### Multi-transcriptome meta-analysis to increase power of expression QTLs detection

Based on shared genetic effects of all types of expression QTLs across tissues, a multi-transcriptome meta-analysis was introduced to increase the power to detect sQTLs, eeQTLs and geQTLs (Figure 6, Table 3). For sQTLs, eeQTLs and geQTLs that had significant effects (p < 0.05) in all of white blood cells, milk cells and muscle transcriptomes, their standardised effects (signed t values) in each transcriptome were simply combined and tested for significance against a χ^2^ distribution with 1 degree of freedom. Overall, the multi-transcriptome meta-analysis based on summary statistics substantially increased the power of expression QTLs detection (Figure 6). The significance of multi-transcriptome expression QTLs was compared with their significance in the liver transcriptome (Figure 6, Table 3). For a criteria where the expression QTLs had both multi-transcriptome meta-analysis p < 1×10^−5^ and liver transcriptome analysis p < 0.05, all types of expression QTLs had significant overlap of the SNPs between the meta-analysis and the single transcriptome analysis in liver. In fact most of the significant sQTLs, eeQTLs and geQTLs detected by the meta-analysis were also detected by the liver analysis but at a much higher p-value (Table 3).

**Table 3.** The overlap between the multi-transcriptome meta-analysis of three tissues and the single transcriptome results of liver. sQTLs: splicing quantitative trait loci; eeQTLs: exon expression QTLs; geQTLs: gene expression QTLs. Liver: QTLs with single-transcriptome effects p < 0.05 in the liver. Meta-analysis: QTLs with χ^2^ p < 1e-05 for the meta-analysis of white blood cells, milk cells and muscle. ‘+’ indicates the number of SNPs met the significance criteria while ‘-’ indicates number of SNPs failed to meet the significance criteria.

| sQTLs |  | Meta-analysis |  | Overlap p |
| --- | --- | --- | --- | --- |
|  |  | + | - |  |
| Liver | + | 768,650 | 2,842,552 | $p < 2e-16$ |
|  | - | 139,551 | 4,301,536 |  |

| eeQTLs |  | Meta-analysis |  | Overlap p |
| --- | --- | --- | --- | --- |
|  |  | + | - |  |
| Liver | + | 1,052,041 | 3,186,839 | $p < 2e-16$ |
|  | - | 140,033 | 3,575,617 |  |

| geQTLs |  | Meta-analysis |  | Overlap p |
| --- | --- | --- | --- | --- |
|  |  | + | - |  |
| Liver | + | 178,093 | 473,232 | $p < 2e-16$ |
|  | - | 43,578 | 6,663,855 |  |

**Figure 6.**
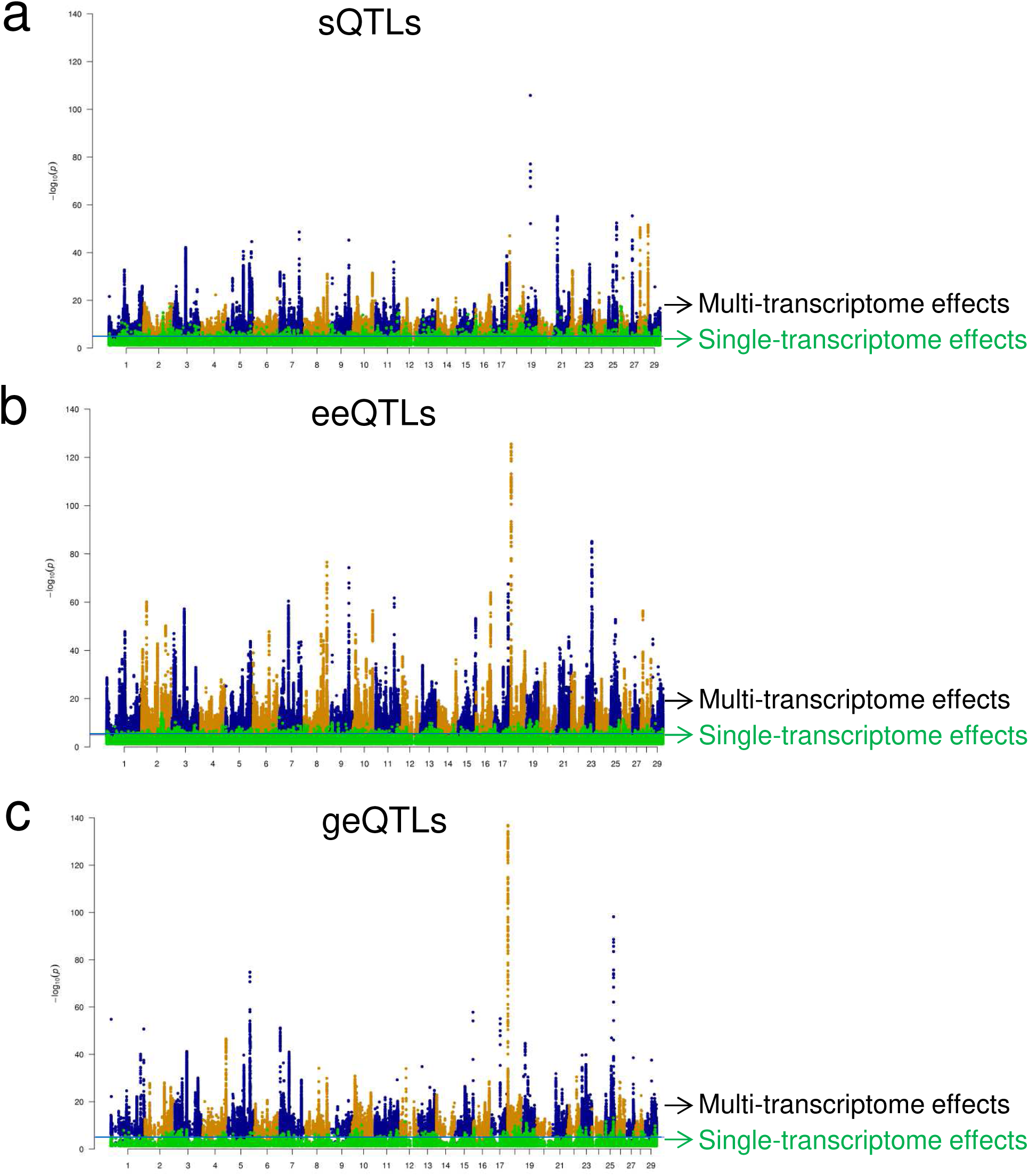
Multi-transcriptome meta-analysis (blood, milk and muscle) for cis splicing sQTLs (**a**), exon expression eeQTLs (**b**) and gene expression geQTLs (**c**). In each panel, the significance of multi-transcriptome effects were tested against a ?2 with 1 degree of freedom for combined expression QTLs effects (dots in blue and orange). These multi-transcriptome effects were shown together with the single-transcriptome effects in liver of the same expression QTLs (dots in green).

### Overlap between expression QTLs and QTL for dairy and beef traits

We examined whether cis sQTLs, eeQTLs and geQTLs were significantly enriched amongst SNPs associated with economically important cattle traits. Pleiotropic SNPs significantly (FDR<0.01) associated with more than one of 24 dairy traits [32] and with more than one of 16 beef traits [33] were tested for overlap with detected sQTLs, eeQTLs and geQTLs (Figure 7, Additional file 10,11: Supplementary Table S8,9). Overall, sQTLs, geQTLs and eeQTLs identified in white blood and milk cells had greater overlap with SNPs associated with dairy and beef traits than sQTLs, geQTLs and eeQTLs identified in liver and muscle (Figure 7). sQTLs in white blood and milk cells were significantly enriched for dairy cattle pleiotropic SNPs, including SNPs from the *CSN3* loci on chromosome 6 (Figure 7a,b). eeQTLs in the white blood cells had the largest absolute amount of SNPs overlapping with dairy cattle pleiotropic SNPs (Figure 7b) and was the only expression QTL type with significant enrichment with beef cattle pleiotropic SNPs (Figure 7b). eeQTLs in milk cells and liver also had significant enrichment for dairy cattle pleiotropic SNPs (Figure 7a). Of the geQTLs, only those from white blood cells had a significant enrichment with dairy cattle pleiotropic SNPs (Figure 7a).

**Figure 7.**
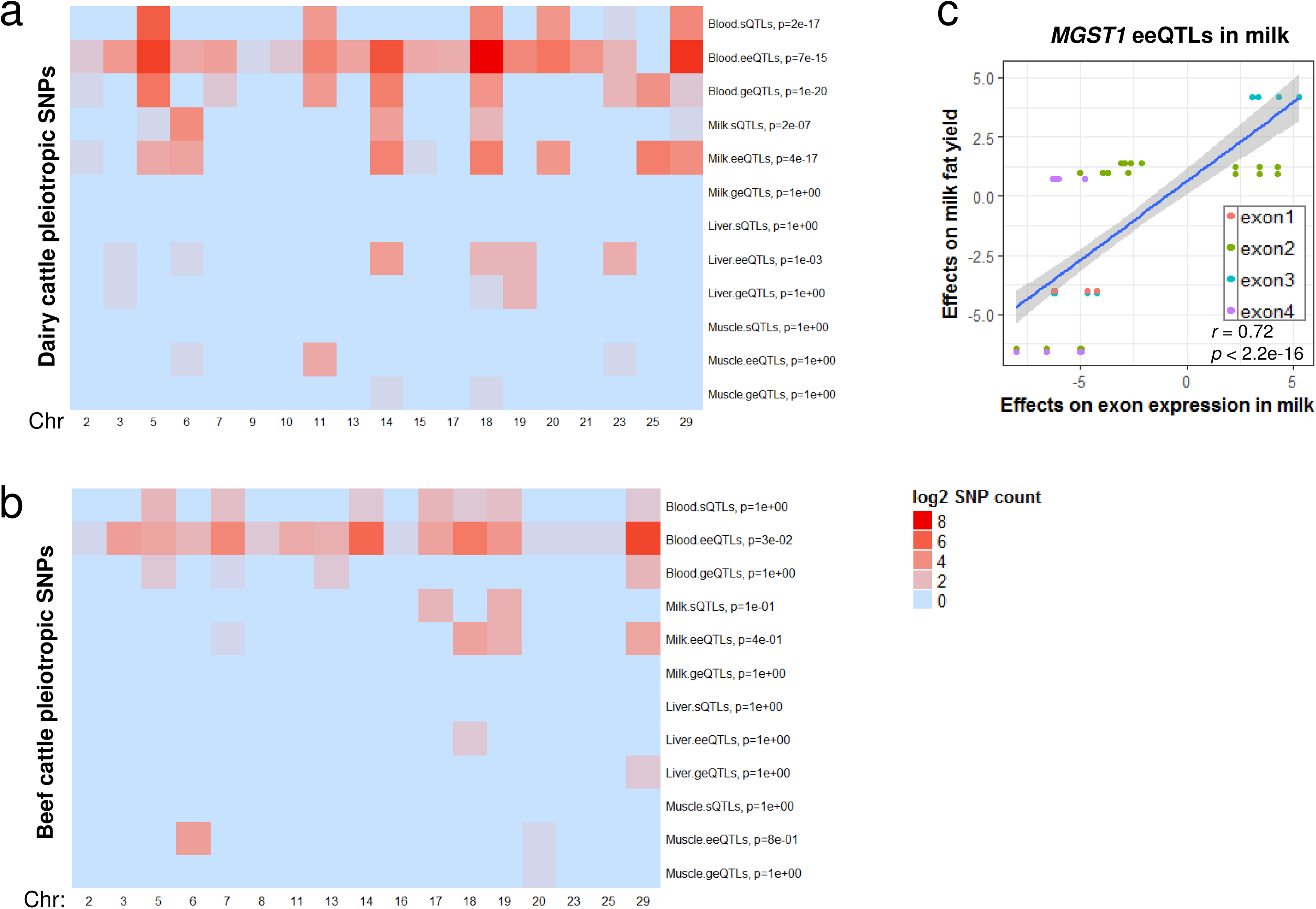
Significance of the overlap, based on the Fisher’s exact test, between pleiotropic QTL for a range of traits in cattle for dairy(**a)** and beef (**b**) and cis splicing quantitative trait loci (sQTLs), exon expression QTL (eeQTLs) and gene expression QTL (geQTLs) in all tissues, where the colour represents the significance of the overlap. Where blood refers to white blood cells and milk refers to milk cells. Significance of the overlap was based on the Fisher’s exact test. Only chromosomes containing overlapping SNPs are shown. **c**) An example of *MGST1* showing the relationship between QTL effects on exon expression in milk cells and their effects on dairy cattle milk fat yield.

An example of an eeQTLs that overlaps a milk production QTL is for *MGST1*, where effects of milk cell eeQTLs were highly significantly associated with their effects on milk fat yield [32] (Figure 7c). Specifically, some expression QTLs with strong associations with the variation in milk cell expression levels of exon 2 (Chr5:93,942,195-93,942,055) and exon 3 (Chr5: 93,939,244-93,939,150) of *MGST1* (Additional file 2: Supplementary Figure S7) also had strong associations with the variation in milk fat yield (Figure 7c). Littlejohn et al.[31] identified SNPs associated with milk yield percentage and *MGST1* expression in the mammary gland, including 17 putative causal variants. Most SNPs identified by Littlejohn et al. originated from whole genome sequence and so were not present on the high density SNP chip data we analysed for dairy cattle pleiotropy [32] (Additional file 12: Supplementary Table S10). However, 53 significant milk cell eeQTLs identified by the current study overlapped with the top 200 SNPs from Littlejohn et al [31] (Additional file 12: Supplementary Table S10), which was significantly more than expected by random chance. The 53 eeQTLs included the SNP suggested as a putative causal candidate (Chr5:93945738) [31], which was significantly associated with the variation in expression level of the third exon (5:93939150-93939244) of *MGST1* in milk cells (Additional file 12: Supplementary Table S10). No milk cell geQTLs was called for *MGST1*, as all of them had weak effects on the whole *MGST1* gene expression in milk cells, resulting in a large FDR (Additional file 12: Supplementary Table S10).

## Discussion

We performed a systematic analysis of cis expression QTLs (<=1Mb) in multiple tissues centred around RNA splicing events, using a large number of RNA and whole genome sequence data from an important domestic animal species. Overall, differential splicing between tissues is ubiquitous and between breeds is common. Differential splicing between individuals due to SNPs (sQTLs) occurs for many genes and is enriched with cattle complex trait QTL. Within each tissue, all cis expression QTLs types showed significant overlap. Most geQTLs and sQTLs were detected as eeQTLs indicating that the exon expression can be altered by changing the expression of the whole gene or by changing the splicing. However, an sQTL was likely to be an eeQTLs and, to a lesser extent, geQTLs. Between tissues, while all QTLs types showed significant overlap between white blood cells and milk cells and between liver and muscle, the strongest cross-tissue sharing appeared to be at the exon level (sQTLs and eeQTLs). This is supported by many significant tissue pair genetic correlations. Such cross-tissue expression QTL sharing allowed the multi-transcriptome meta-analysis of expression QTL effects which substantially increases power to detect significant expression QTLs.

The majority of significant sQTLs were detected from white blood and milk cells (Figure 2a,b) which also overlapped with SNP chip based complex trait QTL (Figure 7), compared to sQTLs detected from liver and muscle. This is probably due to the larger sample size for white blood and milk cells than for liver and muscle (Table 1) and the higher sequencing depth (Additional file 1: Supplementary Methods). One of the significant white blood cell sQTLs (29:44585782) for *CAPN1* is also a SNP chip based significant pleiotropic SNP for 16 beef cattle traits (Additional file 11: Supplementary Table S9). This SNP is associated with shear force in multiple taurine breeds [34].

In the milk cell transcriptome, a significant sQTL (Chr6:87392580, Figure 2a) with predicted splicing function [28] within the fifth exon (6:87,392,578-87,392,750) of *CSN3* is strongly associated with differential splicing between Holstein and Jerseys (Figure 1b). Variants within *CSN3* have long been found to be associated with milk traits [35, 36] but only recently have putative causal variants been prioritised [26]. The milk cell sQTL 6:87392580 had perfect linkage disequilibrium (r =1) with the variant 6:87390576 which has been suggested as a putative causal variant for effects on milk protein yield and percentage [26, 27]. Given it is at a splicing site, 6:87392580 could be a putative causal variant contributing to milk production in dairy cattle by altering exon splicing.

Compared to identified bovine cis geQTLs, cis sQTLs tended to be closer to the transcription starting site (TSS) and had highest concentrations of intronic SNPs (Figure 3). In humans, cis sQTLs [9, 37] were more enriched for intron SNPs than other types of QTLs. However, reports of the distance between human QTLs and TSS appear to be inconsistent. While no difference in enrichment of SNPs near TSS between sQTLs and geQTLs were found by the human GTEx project [8], a more recent study[9] found that human geQTLs were more enriched near TSS than sQTLs. Our results appear to stand in between the results of GTEx project and the later findings from Li et. al. [9], where cattle sQTLs were slightly closer to TSS than geQTLs. However, this difference is not significant in all tissues (Figure 3a). On the other hand, significant overlap between sQTLs and geQTLs was found in this study (Figure 4) and by the human GTEx project [8]. However, Li et. al. [9] found that human cis sQTLs were independent of geQTLs. These inconsistent observations are likely to be due to a number of differences between these studies, including definition of sQTLs, choice of tissues and populations and computational procedures. Also, these inconsistent observations also suggest that we are still at the very early stage of understanding of sQTLs features.

Within each studied bovine tissue, the largest amount of overlap between expression QTL types was found either between exon expression eeQTLs and sQTLs or between eeQTLs and geQTLs (Figure 4). Further, the largest amount of enrichments of cattle pleiotropic SNPs was found for eeQTLs, followed by sQTLs and geQTLs. The white blood cell eeQTLs showed particularly strong enrichments of pleiotropic SNPs for dairy and beef cattle. In a large scale human blood cell expression QTLs study [12], eeQTLs also showed the strongest enrichments of GWAS variants, followed by sQTLs and geQTLs. Thus, focusing on exon-level QTLs, including eeQTLs and sQTLs, could increase the chance of finding regulatory variants for complex traits, as proposed by Guan et. al.[38].

A hypothesis to explain these results is that mutations in regulatory DNA may increase the expression of one or more transcripts from a gene. If they increase expression of one transcript then they may be detected as an eeQTL for the exons in that transcript, as a sQTL for exons spliced out of that transcript or as a geQTL if this transcript forms a large part of the total transcription from the gene. Thus, there is expected to be overlap between eeQTLs, geQTLs and sQTLs, but at least sQTLs and eeQTLs should overlap and this is what we found (Figure 4, Additional file 9: Supplementary Table S7). It appears that eeQTLs detect the largest proportion of these regulatory polymorphisms provided sequencing depth is high.

In humans, significant cross-tissue sharing of sQTLs and geQTLs was reported [8, 39]. In our study of cattle, the strongest evidence of expression QTL sharing appeared to be at the exon level. This included sQTLs and eeQTLs sharing between white blood and milk cells and between liver and muscle (Figure 5). When extending the examination of expression QTLs to include those with p < 0.05 (Additional file 2: Supplementary Figure S6), the exon-level expression QTLs cross-tissue sharing is also the greatest.

We highlighted a few examples of cross-tissue shared eeQTLs along with the related exons, of which the genetic correlations of the expression and splicing in different tissues were significant (Figure 5c). One of these eeQTLs is located within the milk fat yield [27, 31] QTL *MGST1* (Figure 5c, Additional file 2: Supplementary Figure S7). For eeQTLs associated with *MGST1*, a strong positive relationship of SNP effects was observed between milk cell eeQTLs and dairy milk fat yield SNPs (Figure 7c). Furthermore, the identified milk cell eeQTL overlaps with previously identified putative causal variants [31] within *MGST1* for milk fat percentage, thus supporting their candidacy. This overlap further supports the top candidate SNP 5:93945738 with significant effects on the abundance of the third exon of *MGST1* (Additional file 12: Supplementary Table S10) for milk fat traits. Overall, our analysis demonstrates the significant potential of using detailed exon analysis to aid in identification of putative causative mutations.

Based on the sharing of expression QTLs between tissues, a multi-transcriptome meta-analysis which simply combined expression QTL effects to substantially increase the power (Figure 6) was introduced. Using this approach, combined expression QTL effects of white blood cells, milk cells and muscle were validated in the liver (Table 3). This also demonstrated the significant extent of QTL sharing across tissues. Previously, Flutre et. al. [39] combined data from human fibroblasts, lymphoblastoid cell lines and T-cells and found that up to 88% of geQTLs were shared across tissues at FDR<0.05 level. We checked the existing results of the meta-analysis combining SNP effects from tissues of white blood cells, milk cells and muscle at the FDR threshold < 0.05. We found that the meta-analysis identified 585,406 geQTLs with FDR < 0.05 in more than one tissue. This accounted for 69.2% of total geQTLs (845,431) that were called and common in the individual geQTL analysis of white blood cells, milk cells and muscle. While there were differences in the selection of tissue/cell type between our experiment and Flutre et. al, it is possible that the analysis proposed by Flutre et. al with more complex procedures would be more powerful than the meta-analysis introduced by us. Flutre et. al applied principal components analysis to normalise their gene expression data while we used quantile normalisation which appeared to show good performances in combining different transcriptome datasets [40]. However, our meta-analysis is powerful for detecting and validating many expression QTLs that have an effect in the same direction in multiple tissues, and is simpler to implement than that of Flutre et al. A future systematic comparison of different approaches of analysing expression QTL in multiple tissues would be very useful.

As one of earliest investigations of large animal expression QTLs, our study has its potential limitations. While the overlaps between sQTLs detected with exon and intron analyses were significantly more than expected by random chance, the absolute amount of overlap was still small. Through all analyses, there were always many more splicing events detected by intron analyses implemented by leafcutter [10] than the exon analysis (Additional file 5: Supplementary Table S3). This appears to be consistent with Li et al [10], the authors of leafcutter. They suggested that intron-centred analyses can be both more sensitive (lower proportion of false negatives) and more accurate (lower proportion of false positives) than the exon based splicing mapping methods, such as Altrans [41].

We found that the strongest sharing of expression QTLs was either between white blood and milk cells or between liver and muscle tissues, at the threshold of FDR <0.01 (Figure 5). The white blood and milk cells sampled from the same Holstein and/or Jersey cattle of experiment III had a larger sample size and higher read coverage, compared to the liver and muscle tissues sampled from different Angus bulls of experiment IV. The reduced expression QTL sharing detected between, e.g., muscle and milk cells, could be due to differences in the tissue, the physiological state of the cattle or the breed. However, it can be also due to different power in the milk cells, liver and muscle datasets compared to the white blood cell data. Nevertheless, in the multi-transcriptome meta-analysis where expression QTLs with low threshold were examined (p<0.05), the combined effects of all types of expression QTLs of the three tissues from different experiments were highly significant (Figure 6). Many of these expression QTLs were also found in liver with p<0.05 (Table 3). This evidence supports the proposal that the sharing of cis expression QTL is extensive across tissues, but these shared expression QTLs may not necessarily have strong effects in each studied tissue. In the latest human expression QTLs mapping study (GTEx consortium) where RNA seq data of 44 tissues from up to 450 individuals were analysed, cis expression QTL tended to be either shared across most tissues or specific to a small subset of tissues [11]. As sample numbers for each tissue increased, GTEx consortium identified more tissue specific expression QTLs [11]. Future studies with significantly increased power and selection of cattle tissues and breeds may update our current results.

Another potential limitation of our study is the use of imputed sequence data, which may introduce imputation errors that lead to inaccurate identification or exclusion of expression QTLs. However, the average imputation accuracy of the 1000 bull genome project data used in this study was high (0.926) [21] and there was a good consistency between the imputed sequence genotypes and RNA sequence genotypes (average concordance = 0.943, Additional File 2: Supplementary Figure S3). Stringent thresholds were also imposed to control the false discovery rate of expression QTLs mapping (either FDR<0.01 or FDR approximately < 0.01 for sQTLs). In the current study, we did not consider the case where a haplotype can be potentially associated with expression phenotypes. While a haplotype analysis can be informative, it would require a very large sample size to achieve reliable results due to testing a large number of combinations of haplotype blocks. In a human study where over 2,000 individuals were analysed, expression QTLs conditioning on expression levels of transcription factor genes were reported [12]. Finally, our results obtained from genome-wide associations do not necessarily contain causal relationships. However, our findings are important for prioritising informative SNP candidates for future validation of causal relationships.

## Conclusions

We found that eeQTLs overlapped with both geQTLs, due to polymorphisms affecting the level of expression of the whole gene, and with sQTLs, due to polymorphisms affecting the exon usage within the gene. sQTLs tended to be closer to the transcription start sites more often located in introns than geQTLs. We found the largest number of sQTLs in white blood cells probably because the power to find them was greatest in this dataset. However, many of the sQTLs found in other tissues were also detected in blood cells and many sQTLs found in blood could be detected in other tissues at higher p-values. The genetic correlation between expression QTLs in different tissues was often indistinguishable from 1.0 indicating that many expression QTLs operate in a similar way across tissues. Consequently, combining results from several tissues using the multi-transcriptome meta-analysis increased power to detect all 3 types of expression QTLs. The potential of exon-level QTLs information was demonstrated by the identification of several strong candidates of putative causal mutations for complex traits: sQTL 6:87392580 within *CSN3* for milk production and eeQTL 5:93945738 within *MGST1* for milk fat yield.

## Methods

### Sample collection

For Experiment I, the sampling of 18 tissues from one lactating Holstein cow followed procedures described by Chamberlain et al [24]. For Experiment II and III, the sampling and processing of all tissues including white blood and milk cells is detailed in Additional file 1: Supplementary Methods. Briefly, animals of Experiment II and III were selected from Agriculture Victoria Research dairy herd at Ellinbank, Victoria, Australia. In Experiment II, milk and mammary tissue samples were taken from six Holstein cows. In the Experiment III, milk and blood samples were originally taken from 112 Holstein and 29 Jersey cows, but only RNA sequence data of 105 Holstein and 26 Jersey with > 50 million reads for milk cells or >25 million reads for white blood cells and had aconcordant alignment rate [18] >80% were used. For Experiment IV, the sampling of 41 *semitendinosus* muscle and 35 liver from Angus bulls was previously described by [42, 43]. As recommended by ENCODE guidelines (https://www.encodeproject.org/about/experiment-guidelines/) biological replicates were favoured over technical replicates for experiments II-IV. However Chamberlain et al [24] assessed technical replicates for experiment I.

### RNA seq data

For Experiment I, RNA extraction and sequencing followed the procedures described by Chamberlain et al [24]. For Experiment II and III, the RNA extraction and sequencing procedure is detailed in Additional file 1: Supplementary Methods. For Experiment IV, RNA extraction and sequencing is previously described by Khansefid et al. [43]. For all experiments sequence quality was checked and were aligned to the Ensembl UMD3.1 bovine genome assembly using TopHat2 [18]. The RNA sequence data processing and quality checking are detailed in Additional file 1: Supplementary Methods.

### Whole genome seq data

Experiments III and IV had whole genome sequence genotypes imputed from the SNP chip genotypes using FImpute [44] based on the 1000 bull genomes project [19]. The overall imputation accuracy of the most recent genome sequence data ranged from 0.898 to 0.952 depending on chromosomes [21]. 50K Illumina genotypes were used for imputation for experiment III with previous protocols [27, 45]. For experiment IV, 800K and 50K Illumina genotypes were used with procedures following [33]. SNPs were filtered for minor allele frequency > 0.01 and resulted in 14,302,604 and 13,632,145 SNPs used in the analysis in experiment III and IV, respectively. There were 10,242,837 SNPs shared between experiment III and IV.

### Gene/exon analysis

Gene count data were generated by Python package HTSeq [46] using default settings. The exon count data were generated by Bioconductor package featureCounts [47] in R v3.3.2 [48]. The Ensembl based bovine genome reference (UMD3.1) was used to define genes and exons. Genes and exons with count per million >0 in more than 40% of RNA samples were used for all the following analyses. This filtering allowed the analysis to focus on exons or genes with relatively robust expression in many RNA sequencing samples. The exon-based tissue principal components analysis used DEseq2 based on the 250 exons, the expression of which were most variable across studied tissues [49]. The usage of 500 and 1,000 exons with the most variable expression across tissue samples were also tested. Consistent with [49], the selection of different numbers of exons had little impact on the clustering patterns (Supplementary Figure S4). The significance of the clustering was determined using ellipse method proposed by [22] and implemented in ggplot2 [50]. The confidence interval was set to 0.95 to which ellipses were drawn based on data categorised by tissue types or by experiments. The separation of ellipses indicated independence of categories of data. The phenotype of exon inclusion was calculated as the exon to gene expression ratio. The phenotype of intron excision was estimated using the publically available software leafcutter [9, 10]. Briefly, leafcutter used RNA seq BAM files as input and generated ratios of reads supporting each alternatively excised intron as the intron excision phenotype [10](http://davidaknowles.github.io/leafcutter/). Those exons and introns with ratio values <0.001 were removed and the remaining ratio values were transformed to log_2_scale, then underwent exon/intron –wise quantile normalisation and individual-wise zscore standardisation [51].

### Gene differential splicing

Both exon inclusions and intron excisions were analysed and used in combination for gene differential splicing for (1) the overall tissue effects and (2) the breed effects. Primarily, differential splicing was defined for the gene containing exons whose variation in inclusion ratios were significantly (FDR<0.1) associated with the tissue or breed variable. To be called as significantly spliced exons, they were required to have at least one adjacent intron whose variation in excision ratios were also significantly (FDR<0.1) associated with the tissue or breed variable. The tissue effects were analysed in a linear mixed model in lme4 [52] in R as:

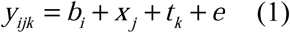

Where *y* = exon inclusion or intron excision ratios, *b*_*i*_ = the animal random effects (*i*=214), *x*_*j*_= the experiments (*j*=4), *t*_*k*_ = tissue type (*k*=19), *e*= random residual term. The fitting of the animal random effects accounted for the fact that only 1 animal was used in experiment I. The P values of F tests were calculated using Satterthwaite approximation implemented in lmerTest [53]. The breed effects for the milk transcriptome data were analysed in a linear model in R as:

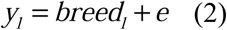

Where *y* = exon inclusion or intron excision ratios in the milk cell transcriptome, *breed*_*l*_ = breeds (l=2, Holstein and Jersey). p values of exons/introns for the tissue effects in equation (1) or the breed effects in equation (2) were used to calculate the false discovery rates (FDR) using qvalue [54] in R. The FDR threshold of such detected exon/intron group was considered as approximately 0.1× 0.1 = 0.01, as a combination of FDR thresholds of exon and intron analyses to reflect our selection criteria for significant splicing events. For genes showing significant differential splicing for (1) the overall tissue effects and (2) the breed effects as described above, enrichments of biological pathway were tested using GOrilla [55]. As many genes had differential splicing events associated with tissue differences, top 10% of the genes with significant differential splicing were selected based on the approximate FDR with combined FDR values of both exon and intron analyses.

### cis expression splicing QTLs

Only transcriptomic data of experiment III and IV were used in sQTLs mapping. Similar to differential splicing analysis described above, a significant (FDR<0.01) cis splicing QTLs was expected to satisfy two conditions simultaneously: (1) a SNP, within or up to ± 1Mb away from the exon, was significantly (FDR<0.1) associated with the variation in the exon inclusion ratio and (2) the same SNP was significantly (FDR<0.1) associated with at least one event of the excision of the intron next to the same exon at the same significance level. Both individual exon inclusion and intron excision values were used as phenotype to map associated QTLs with widely used [8] Matrix eQTL[56] package in R. For each cell type of the experiment III (white blood and milk cells) and experiment IV (liver and muscle), SNPs ± 1Mb from the exon or intron were tested for regressions with the exon inclusion or intron excision phenotype. For milk cell transcriptome, breed was fitted as a covariate.

To compare cis sQTLs with exon expression cis eeQTLs and cis gene expression geQTLs, the expression count data were normalised by voom [57] estimating mean-variance relationship to calculate observation-level weighted expression values. Normalised expression values of exons and genes were used as phenotype to map cis expression QTLs (within ± 1Mb) at FDR <0.01 level as described above.

### SNP annotation

The gene transcription start site coordinates were downloaded from Ensembl (http://www.ensembl.org) and the absolute difference between the position of a SNP and the transcription start site of the gene were calculated for the SNP with significant cis effects. The SNP functional categories were generated using predictions from Ensembl Variant Effect Predictor [28] in conjunction with NGS-SNP [29]. All analysed SNPs were assigned a functional category.

### Dairy and beef cattle pleiotropic QTL

To test the significance of overlap between cis expression QTLs and SNPs associated with cattle phenotype, meta-analyses of dairy and beef cattle pleiotropy were performed using single-trait GWAS results from Xiang et al [32] and Bolormaa et al [33]. HD 800K SNP chip genotypes were used for trait GWAS. 24 dairy cattle traits with matching phenotype in 9,662 bulls and cows and 16 beef cattle traits with animal numbers >2,000 were selected. Briefly, the multi-trait χ^2^ statistic for the *i*th SNP was calculated based on its signed t values generated from each single trait GWAS [33]:

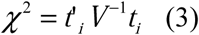

For dairy cattle, the meta-analysis was based on the weighted SNP effects *t*_*w*_ combining SNP effects calculated separately in bulls and cows. The *t*_*w*_ accounting for the phenotypic error differences between bulls and cows [27] was calculated as:

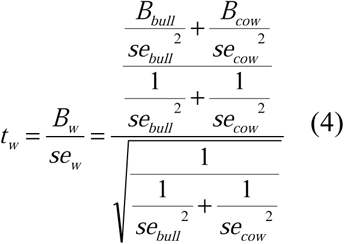

Where the weighted SNP t value *t*_*w*_ was the quotient of the weighted SNP effects *B*_*w*_ and the weighted effect error *se*_*w*_. *B*_*bull*_ and *se*_*bull*_ were the SNP effects and error obtained from single-trait GWAS in bulls and *B*_*cow*_ and *se*_*cow*_ were the SNP effects and error of cows. Those SNPs which had meta-analysis FDR<0.01 were chosen to be compared with cis expression QTLs. The lead SNP loci were defined as ±1 Mb from the lead SNPs identified in the previous analysis [32] and [33].

### The significance of overlaps

The significance of overlaps were compared with the expected number using the Fisher’s exact test (*p*) implemented in GeneOverlap [58] in R. This analysis required four types of counts: the size of overlap between set A (e.g., SNPs that were blood sQTLs) and set B (e.g., SNPs that were milk sQTLs), the size of set A, the size of set B and the size of background. The union number of whole genome sequence SNPs with MAF>0.01 in each breed and the bovine high density chip SNPs were used as the background. Where expression QTL categories from different breeds of dairy and beef cattle were tested for overlap, the number of common SNPs between breeds was used.

### Genetic correlations using local genomic relationship matrices

The cross-tissue sharing of SNPs were confirmed by bivariate GREML analysis using GCTA [59]. For an exon or a gene of interest, its inclusion ratios or expression levels in two different tissues were treated as two different phenotype, *tr*_*1*_ and *tr*_*2*_. The SNPs within 1Mb of this exon or gene were used to make a local genomic relationship matrix, i.e., LGRM, representing the local polygenic component *a* with potential associations with the variation in the splicing or expression level of the exon or gene. This allowed linear mixed modelling of the local additive genetic variances of *tr*_*1*_, var_lg_(*tr*_*1*_) and of *tr*_*2*_, var_lg_(*tr*_*2*_) and the local additive genetic covariance between *t*_1_ and *t*_2_, cov_lg_(*tr*_*1*_,*tr*_*2*_) using GREML [59]. This approach agreed with the definition of *cis* expression QTLs defined in this study (also within 1Mb distance to the exon or gene) and allowed the estimation of genetic correlation 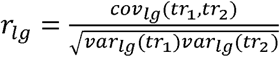 (5). Genetic correlations were also tested for their significance of being different from 0 or 1, by fixing the correlation value to 0 and 1 using GCTA [59].

### Validation by multi-transcriptome meta-analysis

The validation based on expression QTL effect commonality across tissues was conducted by comparing the combined expression QTL effects from white blood cells (experiment III), milk cells (experiment III) and muscle (experiment IV) transcriptomes with their effects in the liver transcriptome (experiment IV). The standardised expression QTL effects, b/se, signed t values were calculated from single-transcriptome results of white blood cells (*t*_*1*_), milk cells (*t*_*2*_) and muscle (*t*_*3*_). The significance of multi-transcriptome effects of an expression QTL was tested by χ^2^ distribution with 1 degree of freedom:

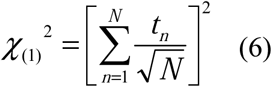

*N* = the number of studied tissues (*N*=3 in this case) where the original SNP t values were estimated. Provided the individual t-values followed a t-distribution under the null hypothesis, the properties of the average t value in the current study was a simple mathematical result which approximated the chi square distribution with 1 degree of freedom, the null hypothesis of which was that the SNP does not have any significant associations in any of the 3 tissue types. Previously, the concept of meta-analysis combining SNP t values estimated from different datasets has been also applied to analyse multiple quantitative phenotypic traits in large animals to increase power (see [33, 60] and equation (3)). The expression QTLs that participated in the validation analysis had single-transcriptome effect p < 0.05 in each tissue and the significance of the multi-transcriptome effects was defined as p < 1e-05. Significant multi-transcriptome expression QTLs were compared with the liver single-transcriptome effects at p < 0.05 level. We chose to combine two tissues which appeared to display strong power (white blood and milk cells, experiment III) with the third tissue from a different experiment with relatively weak power (muscle, experiment IV). The single tissue left to be compared with was liver, a tissue which also appeared to show weak power and was from experiment IV. These choices intended to create enough differences for the meta-analysis to combine the SNP effects and for the combined SNP effects to be compared with the SNP effects in the single tissue.

## List of abbreviations

sQTLs: splicing quantitative trait loci
geQTLs: gene expression quantitative trait loci
eeQTLs: exon expression quantitative trait loci

## Declarations

### Ethics approval and consent to participate

For experiment I-III, the animal ethics was approved by the Victoria Animal Ethics Committee (application number 2013-14), Australia. For experiments IV, the animal ethics was approved by the University of New England Animal Ethics Committee (AEC 06/123, NSW, Australia) and Orange Agriculture Institute Ethics Committee (ORA09/015, NSW, Australia).

### Consent for publication

Not applicable.

### Availability of data and materials

The RNA sequence data for experiment I was published [24] (NCBI Sequence Read Archive, SRA, accession SRP042639); For experiment II: SRA accession SRP111067; For experiment III: SRA accession PRJNA305942; For experiment IV: SRA accession PRJNA392196; The imputed whole genome sequence data was part of the 1000 bull genome project [19].

## Author contributions

A.J.C., R.X., M.E.G., B.J.H and H.D.D. conceived the experiments. C.P.P., C.M.R., B.A.M., J.B.G., L.C.M., Y.C. and A.J.C. performed sample collections and RNA sequencing experiments. S.B., I.M.M., M.K. and H.D.D. provided data and assisted with study design. R.X., B.J.H., C.J.V., A.J.C., M.E.G, P.J.B and Z.Y. analysed data. R.X. and M.E.G. wrote the paper. R.X., M.E.G., B.J.H, A.J.C., I.M.M and H.D.D.. revised the paper. All authors read and approved the final manuscript.

## Acknowledgements

We thank DataGene for access to data used in this study and Gert Nieuwhof, Kon Konstantinov and Timothy P. Hancock for preparation and provision of data.

## Additional files

Additional file 1 (DOCX): Supplementary Methods.

Additional file 2 (PDF): Supplementary Figure S1-7.

Additional file 3 (XLSX): Supplementary Table S1. RNA-seq reads mapped to different bovine genome origins.

Additional file 4 (XLSX): Supplementary Table S2. Splicing junction annotation analysis using BAM files generated by different alignment software.

Additional file 5 (XLSX): Supplementary Table S3. 2×2 tables for the overlap between exon and intron analyses.

Additional file 6 (XLSX): Supplementary Table S4. Genes that display significant differential splicing cross tissues.

Additional file 7 (XLSX): Supplementary Table S5. Genes that display significant differential splicing between breeds.

Additional file 8 (XLSX): Supplementary Table S6. Summary of significant cis splicing sQTLs (within 1Mb distance to the exon).

Additional file 9 (XLSX): Supplementary Table S7. Summary table for count values of sQTLs and eeQTLs in white blood and milk cells.

Additional file 10 (XLSX): Supplementary Table S8. SNP overlap between expression QTLs and SNPs with pleiotropic effects on conventional traits of dairy cattle.

Additional file 11 (XLSX): Supplementary Table S9. SNP overlap between expression QTLs and SNPs with pleiotropic effects on conventional traits of beef cattle.

Additional file 12 (XLSX): Supplementary Table S10. SNP overlap between blood eeQTLs and putative causal variants identified by Littlejohn et al (2016).

## Reference

1. Bourneuf E, Otz P, Pausch H, Jagannathan V, Michot P, Grohs C, Piton G, Ammermüller S, Deloche M-C, Fritz S: Rapid Discovery of De Novo Deleterious Mutations in Cattle Enhances the Value of Livestock as Model Species. Sci Rep 2017, 7.

2. Hayes BJ, Lewin HA, Goddard ME: The future of livestock breeding: genomic selection for efficiency, reduced emissions intensity, and adaptation. Trends Genet 2013, 29:206–214.

3. Meuwissen T, Hayes B, Goddard M: Genomic selection: A paradigm shift in animal breeding. Animal frontiers 2016, 6:6–14.

4. Andersson L, Archibald AL, Bottema CD, Brauning R, Burgess SC, Burt DW: Coordinated international action to accelerate genome-to-phenome with FAANG, the Functional Annotation of Animal Genomes project. Genome Biol 2015, 16.

5. Albert FW, Kruglyak L: The role of regulatory variation in complex traits and disease. Nature reviews Genetics 2015, 16:197.

6. Schaub MA, Boyle AP, Kundaje A, Batzoglou S, Snyder M: Linking disease associations with regulatory information in the human genome. Genome Res 2012, 22:1748–1759.

7. Bouwman AC, Daetwyler HD, Chamberlain AJ, Ponce CH, Sargolzaei M, Schenkel FS, Sahana G, Govignon-Gion A, Boitard S, Dolezal M, et al: Meta-analysis of genome-wide association studies for cattle stature identifies common genes that regulate body size in mammals. Nat Genet 2018, 50:362–367.

8. Consortium G: The Genotype-Tissue Expression (GTEx) pilot analysis: Multitissue gene regulation in humans. Science 2015, 348:648–660.

9. Li YI, van de Geijn B, Raj A, Knowles DA, Petti AA, Golan D, Gilad Y, Pritchard JK: RNA splicing is a primary link between genetic variation and disease. Science 2016, 352:600–604.

10. Li YI, Knowles DA, Humphrey J, Barbeira AN, Dickinson SP, Im HK, Pritchard JK: Annotation-free quantification of RNA splicing using LeafCutter. Nat Genet 2018, 50:151.

11. Consortium G: Genetic effects on gene expression across human tissues. Nature 2017, 550:204.

12. Zhernakova DV, Deelen P, Vermaat M, van Iterson M, van Galen M, Arindrarto W, van’t Hof P, Mei H, van Dijk F, Westra H-J: Identification of context-dependent expression quantitative trait loci in whole blood. Nat Genet 2017, 49:139–145.

13. Mazzoni G, Kadarmideen HN: Computational Methods for Quality Check, Preprocessing and Normalization of RNA-Seq Data for Systems Biology and Analysis. In Systems Biology in Animal Production and Health, Vol 2. Springer; 2016: 61–77

14. Okonechnikov K, Conesa A, García-Alcalde F: Qualimap 2: advanced multi-sample quality control for high-throughput sequencing data. Bioinformatics 2015, 32:292–294.

15. Wang L, Wang S, Li W: RSeQC: quality control of RNA-seq experiments. Bioinformatics 2012, 28:2184–2185.

16. Kim D, Langmead B, Salzberg SL: HISAT: a fast spliced aligner with low memory requirements. Nat Methods 2015, 12:357.

17. Dobin A, Davis CA, Schlesinger F, Drenkow J, Zaleski C, Jha S, Batut P, Chaisson M, Gingeras TR: STAR: ultrafast universal RNA-seq aligner. Bioinformatics 2013, 29:15–21.

18. Kim D, Pertea G, Trapnell C, Pimentel H, Kelley R, Salzberg SL: TopHat2: accurate alignment of transcriptomes in the presence of insertions, deletions and gene fusions. Genome Biol 2013, 14:R36.

19. Daetwyler HD, Capitan A, Pausch H, Stothard P, Van Binsbergen R, Brøndum RF, Liao X, Djari A, Rodriguez SC, Grohs C: Whole-genome sequencing of 234 bulls facilitates mapping of monogenic and complex traits in cattle. Nat Genet 2014, 46:858–865.

20. Li H, Handsaker B, Wysoker A, Fennell T, Ruan J, Homer N, Marth G, Abecasis G, Durbin R: The Sequence Alignment/Map format and SAMtools. Bioinformatics 2009, 25:2078–2079.

21. Pausch H, MacLeod IM, Fries R, Emmerling R, Bowman PJ, Daetwyler HD, Goddard ME: Evaluation of the accuracy of imputed sequence variant genotypes and their utility for causal variant detection in cattle. Genet Sel Evol 2017, 49:24.

22. Fox J, Weisberg S: An R companion to applied regression. Sage Publications; 2011.

23. Cánovas A, Rincón G, Bevilacqua C, Islas-Trejo A, Brenaut P, Hovey RC, Boutinaud M, Morgenthaler C, VanKlompenberg MK, Martin P: Comparison of five different RNA sources to examine the lactating bovine mammary gland transcriptome using RNA-Sequencing. Sci Rep 2014, 4.

24. Chamberlain AJ, Vander Jagt CJ, Hayes BJ, Khansefid M, Marett LC, Millen CA, Nguyen TTT, Goddard ME: Extensive variation between tissues in allele specific expression in an outbred mammal. BMC Genomics 2015, 16:993.

25. Li YI, Knowles DA, Pritchard JK: LeafCutter: Annotation-free quantification of RNA splicing. bioRxiv 2016:044107.

26. MacLeod I, Bowman P, Vander Jagt C, Haile-Mariam M, Kemper K, Chamberlain A, Schrooten C, Hayes B, Goddard M: Exploiting biological priors and sequence variants enhances QTL discovery and genomic prediction of complex traits. BMC Genomics 2016, 17:1.

27. Kemper KE, Reich CM, Bowman PJ, vander Jagt CJ, Chamberlain AJ, Mason BA, Hayes BJ, Goddard ME: Improved precision of QTL mapping using a nonlinear Bayesian method in a multi-breed population leads to greater accuracy of across-breed genomic predictions. Genet Sel Evol 2015, 47:1.

28. McLaren W, Gil L, Hunt SE, Riat HS, Ritchie GR, Thormann A, Flicek P, Cunningham F: The Ensembl Variant Effect Predictor. Genome Biol 2016, 17:1.

29. Grant JR, Arantes AS, Liao X, Stothard P: In-depth annotation of SNPs arising from resequencing projects using NGS-SNP. Bioinformatics 2011, 27:2300–2301.

30. National Center for Biotechnology Information (NCBI) [https://www.ncbi.nlm.nih.gov/]

31. Littlejohn MD, Tiplady K, Fink TA, Lehnert K, Lopdell T, Johnson T, Couldrey C, Keehan M, Sherlock RG, Harland C: Sequence-based Association Analysis Reveals an MGST1 eQTL with Pleiotropic Effects on Bovine Milk Composition. Sci Rep 2016, 6.

32. Xiang R, MacLeod I, Bolormaa S, Goddard M: Genome-wide comparative analyses of correlated and uncorrelated phenotypes identify major pleiotropic variants in dairy cattle. Sci Rep 2017, 7:9248.

33. Bolormaa S, Pryce JE, Reverter A, Zhang Y, Barendse W, Kemper K, Tier B, Savin K, Hayes BJ, Goddard ME: A multi-trait, meta-analysis for detecting pleiotropic polymorphisms for stature, fatness and reproduction in beef cattle. PLoS Genet 2014, 10:e1004198.

34. McClure M, Ramey H, Rolf M, McKay S, Decker J, Chapple R, Kim J, Taxis T, Weaber R, Schnabel R: Genome-wide association analysis for quantitative trait loci influencing Warner–Bratzler shear force in five taurine cattle breeds. Anim Genet 2012, 43:662–673.

35. Kühn C, Freyer G, Weikard R, Goldammer T, Schwerin M: Detection of QTL for milk production traits in cattle by application of a specifically developed marker map of BTA6. Anim Genet 1999, 30:333–339.

36. Glantz M, Gustavsson F, Bertelsen HP, Stålhammar H, Lindmark-Månsson H, Paulsson M, Bendixen C, Gregersen VR: Bovine chromosomal regions affecting rheological traits in acid-induced skim milk gels. J Dairy Sci 2015, 98:1273–1285.

37. Takata A, Matsumoto N, Kato T: Genome-wide identification of splicing QTLs in the human brain and their enrichment among schizophrenia-associated loci. Nature Communications 2017, 8:14519.

38. Guan L, Yang Q, Gu M, Chen L, Zhang X: Exon expression QTL (eeQTL) analysis highlights distant genomic variations associated with splicing regulation. Quantitative Biology 2014, 2:71–79.

39. Flutre T, Wen X, Pritchard J, Stephens M: A statistical framework for joint eQTL analysis in multiple tissues. PLoS Genet 2013, 9:e1003486.

40. Thompson JA, Tan J, Greene CS: Cross-platform normalization of microarray and RNA-seq data for machine learning applications. PeerJ 2016, 4:e1621.

41. Ongen H, Dermitzakis ET: Alternative Splicing QTLs in European and African Populations. Am J Hum Genet 2015, 97:567–575.

42. Chen Y, Gondro C, Quinn K, Herd R, Parnell P, Vanselow B: Global gene expression profiling reveals genes expressed differentially in cattle with high and low residual feed intake. Anim Genet 2011, 42:475–490.

43. Khansefid M, Millen CA, Chen Y, Pryce JE, Chamberlain AJ, Vander Jagt CJ, Gondro C, Goddard ME: Gene expression analysis of blood, liver, and muscle in cattle divergently selected for high and low residual feed intake. J Anim Sci 2017, 95:4764–4775.

44. Sargolzaei M, Chesnais JP, Schenkel FS: A new approach for efficient genotype imputation using information from relatives. BMC Genomics 2014, 15.

45. Erbe M, Hayes B, Matukumalli L, Goswami S, Bowman P, Reich C, Mason B, Goddard M: Improving accuracy of genomic predictions within and between dairy cattle breeds with imputed high-density single nucleotide polymorphism panels. J Dairy Sci 2012, 95:4114–4129.

46. Anders S, Pyl PT, Huber W: HTSeq—a Python framework to work with high-throughput sequencing data. Bioinformatics 2015, 31:166–169.

47. Liao Y, Smyth GK, Shi W: featureCounts: an efficient general purpose program for assigning sequence reads to genomic features. Bioinformatics 2013, 30:923–930.

48. Team RC: R: A language and environment for statistical computing. 2013.

49. Love MI, Huber W, Anders S: Moderated estimation of fold change and dispersion for RNA-seq data with DESeq2. Genome Biol 2014, 15:550.

50. Wickham H: ggplot2: elegant graphics for data analysis. Springer; 2016.

51. Degner JF, Pai AA, Pique-Regi R, Veyrieras J-B, Gaffney DJ, Pickrell JK, de Leon S, Michelini K, Lewellen N, Crawford GE: DNase I sensitivity QTLs are a major determinant of human expression variation. Nature 2012, 482:390–394.

52. Bates D, Mächler M, Bolker B, Walker S: Fitting Linear Mixed-Effects Models Using lme4. Journal of Statistical Software 2015, 67:48.

53. Kuznetsova A, Brockhoff PB, Christensen RHB: lmerTest Package: Tests in Linear Mixed Effects Models. Journal of Statistical Software 2017, 82:26.

54. Bass AJ, Dabney A, Robinson D: qvalue: Q-value estimation for false discovery rate control. R package version 2.9.0, http://github.com/jdstorey/qvalue., vol. 2017; 2015.

55. Eden E, Navon R, Steinfeld I, Lipson D, Yakhini Z: GOrilla: a tool for discovery and visualization of enriched GO terms in ranked gene lists. BMC Bioinformatics 2009, 10:48.

56. Shabalin AA: Matrix eQTL: ultra fast eQTL analysis via large matrix operations. Bioinformatics 2012, 28:1353–1358.

57. Law CW, Chen Y, Shi W, Smyth GK: Voom: precision weights unlock linear model analysis tools for RNA-seq read counts. Genome Biol 2014, 15:R29.

58. Shen L: GeneOverlap: An R package to test and visualize gene overlaps. 2014.

59. Yang J, Lee SH, Goddard ME, Visscher PM: GCTA: a tool for genome-wide complex trait analysis. Am J Hum Genet 2011, 88:76–82.

60. Bolormaa S, Hayes BJ, van der Werf JH, Pethick D, Goddard ME, Daetwyler HD: Detailed phenotyping identifies genes with pleiotropic effects on body composition. BMC Genomics 2016, 17:1.

